# RpoN is required for a functional type III secretion system in *Yersinia pseudotuberculosis*

**DOI:** 10.1101/2022.02.11.480049

**Authors:** A.K.M.F. Mahmud, R. Navais, K. Nilsson, R. Choudhury, K. Avican, M. Fallman

## Abstract

Pathogenic bacteria use a broad range of virulence factors to thrive within their host. *Yersinia pseudotuberculosis*, a gram-negative enteropathogen in humans, utilises a Type III secretion system to overcome the host’s innate immune response. A global regulator previously shown to be essential for virulence in *Y. pseudotuberculosis* is RpoN. We show here that a strain lacking RpoN has a severely reduced capacity to secrete Yop effectors. This strain has a substantially reduced expression of genes encoding structural components of the secretion apparatus, the ysc operons, while expression of genes encoding effectors and translocators is less affected. We show that RpoN regulates a complex network, where one part is suggested to contribute to inducing Type III secretion via the enhancer-binding protein GlrR/RpoN regulon. By analogy, during inducement of Type III secretion, RpoN has a positive effect on expression of the sigma factor RpoE, which also is known to act downstream of GlrR. Interestingly, we found putative RpoE binding sites upstream of the *ysc* operons, and also a partial rescue of Yop secretion in the rpoN mutant strain by overexpression of RpoE. Our findings suggest that the RpoN-mediated effect on expression of type III genes involves a sigma factor hierarchy, where RpoN via RpoE contributes to inducing Type III secretion.

## Introduction

*Yersinia pseudotuberculosis* is a food borne pathogen that causes acute gastroenteritis, mesenteric lymphadenitis and diarrhoeal disease ^1^. Pathogenicity relies mainly on a 70-kb virulence plasmid encoding a type III secretion system (T3SS) and the virulence effectors known as *Yersinia* outer proteins (Yops) ^2,3^. This gram-negative bacterium penetrates the intestinal epithelium and invades lymphoid follicles of Peyer’s patches and the cecum ^4,5^. At these sites, abundant in immune cells, this bacterium can survive extracellularly due to its T3SS, which upon intimate contact with host cells deliver Yop effectors into the cytoplasm of the target cells ^6^. By this delivery of virulence effectors into host cells, *Yersinia* interferes with several key mechanisms of the host’s antimicrobial functions, such as phagocytosis and inflammatory responses ^7^. The primary target cells for Yops delivery are neutrophils, macrophages and dendritic cells, which highlights the importance of disarming phagocytic cells upon infection ^8^.

In our recent study on the role of the sigma factor RpoN in *Y. pseudotuberculosis*, we found that an *rpoN* deletion mutant displayed lower expression of T3SS genes and showed clear virulence attenuation in mice ^9^. RpoN or sigma 54 (*σ*^54^) was originally identified in *Escherichia coli* and *Salmonella* as a regulator of genes involved in nitrogen metabolism and assimilation under nitrogen limiting conditions ^10-12^. It has since been shown, in different bacterial species, to regulate a wide range of critical biological functions such as motility, biofilm formation and quorum sensing ^13-15^. There are many studies that implicate RpoN as a regulator of bacterial virulence ^16-18^. Indeed, RpoN has been suggested to control expression of T3SSs in both *Chlamydia trachomatis* and *Pseudomonas aeruginosa* as well as of the type VI secretion system in *Aeromonas hydrophila* ^19-21^. RpoN also regulates T3SS gene expression and plays a vital role in virulence in the plant pathogens *Erwinia amylovora, E. carotovora* and *Pseudomonas syringae* ^22-25^. In these species, RpoN, along with specific enhancer-binding proteins (EBPs), activates the transcription of sigma factor HrpL, the master regulator of T3SS. RpoN-dependent transcription initiation usually requires activation by bacterial EBPs bound at a distal promoter site, an upstream activation sequence ^26,27^. These EBPs use ATP catalysis to remodel the RNA polymerase (RNAP) DNA binding for initiation of transcription ^28^. EBPs contain an AAA+ domain, which is crucial for ATP hydrolysis and coupling with RpoN ^29^. RpoN-dependent promoters are thought to be silent until EBPs stimulate the RpoN-RNA polymerase (RNAP) complex ^30^. Integration host factor (IHF)-mediated DNA looping facilitates contact between the promoter-bound EBP complex and the inactive RNAP-RpoN complex ^31,32^. EBPs are ubiquitous in bacteria, and they can act on different RpoN-dependent promoters under different environmental conditions ^33,34^.

While a direct involvement of RpoN in the regulation of T3SSs and virulence has been suggested for some bacteria, there are still gaps in the understanding of important mechanistic details. In the case of *Y. pseudotuberculosis*, this includes the regulatory events affecting expression of plasmid genes and, thereby, the mechanism by which this sigma factor can control expression of T3SS genes. In this study, we addressed the regulatory role of RpoN in controlling type III secretion (T3S) in *Y. pseudotuberculosis*. The results obtained show that a strain lacking RpoN displays reduced expression of genes encoding structural T3SS components, the *ysc* operons, with an associated reduced capacity to secrete Yop effectors. We show that RpoN regulates a complex network, where one part is suggested to contribute to inducement of T3S via the enhancer-binding protein GlrR/RpoN regulon. In addition, RpoN has a positive effect on expression of the sigma factor RpoE upon inducing T3S, and the putative ability of RpoE to bind upstream of the *ysc* operons implies a role for this regulator in the inducement of *Y. pseudotuberculosis* T3SS.

## Results

### RpoN is required for a functional T3SS in *Y. pseudotuberculosis*

Data from our recent work ^9^ revealed the downregulation of genes coding for different components of the T3SS in a *rpoN* deletion mutant, Δ*rpoN* (Fig. 1A). It was obvious here that the most pronounced decrease in mRNA abundance was seen for operons encoding for structural components of the secretion apparatus, such as the *yscN-U* and *yscC-L* operons (Fig. 1B). In contrast, abundance of mRNA of genes encoding the translocators YopB and YopD, and many of the effector proteins, was not, or was only slightly affected in the Δ*rpoN* strain (Fig1A). Next, we performed a qPCR analysis to check expression of genes encoding different components of T3SS and validate our RNA-Seq. In support of the RNA-seq data, we observed lower expression of genes *yscC, yscR, yscU* and *yscT*, encoding structural components of the secretion apparatus, whereas there was no significant effect on expression levels of *yopB* and *yopD*, encoding two translocator proteins in the absence of RpoN (Fig. 1C). Given these results, we next examined T3S in this strain. We found that the ∆*rpoN* strain was indeed T3S-defective, showing a clearly reduced amount of Yop effectors in the supernatant (Fig. 1D). Trans-complementation of ∆*rpoN* (Fig. 1D) could restore the wild-type (WT) T3S-positive phenotype. Hence, the defect in T3S observed in ∆*rpoN* probably contributed in a significant manner to the attenuation in virulence as our previous study found ^9^.

**Fig 1.**
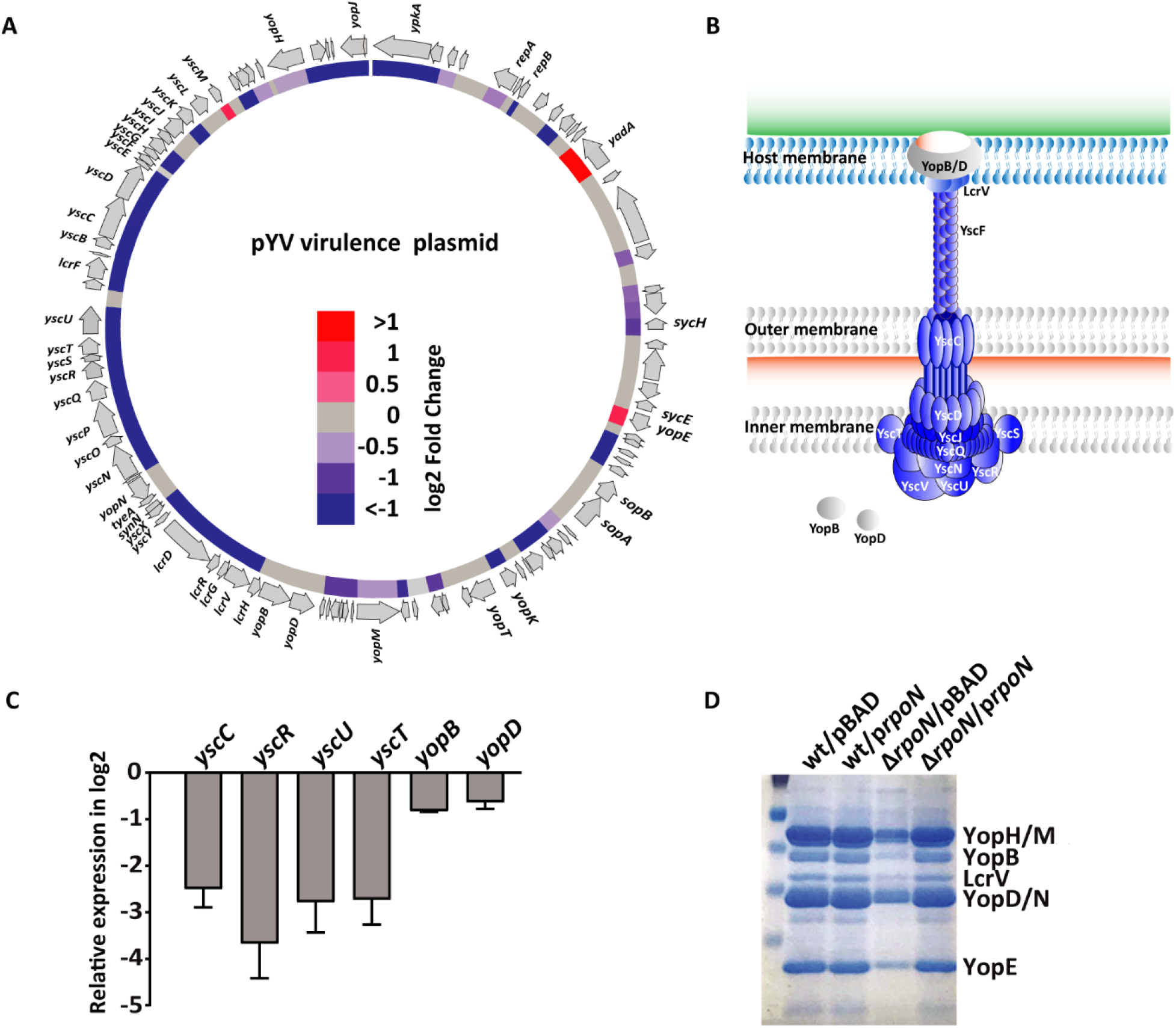
Deletion of *rpoN* in *Y. pseudotuberculosis* leads to reduced expression of virulence plasmid genes encoding T3SS structural components and an associated reduction in secretion of virulence effectors. (A) Schematic representation of the *Yersinia* virulence plasmid (pIBX/pYV), highlighting the genes that were differentially expressed in ∆*rpoN* (P adjusted value <0.05). Inner circle indicates downregulated (blue) and upregulated regions compared to the gene expression in WT. (B) Schematic diagram of the T3SS apparatus. Components whose encoding genes are downregulated in the absence of RpoN are indicated in blue. (C) Relative expression of indicated genes under virulence-inducing conditions in ∆*rpoN* compared with the WT strain. Gene expression was determined by qPCR and represented as relative expression in log2 fold change. Values represent mean ± SEM of three independent experiments. (D) T3S profiles of WT *Y. pseudotuberculosis*, and the isogenic mutant ∆*rpoN* and Δ*rpoN* complemented in *trans* with the pBAD vector expressing RpoN (Δ*rpoN*/p*rpoN*). The WT and Δ*rpoN* strains carried the empty pBAD vector. T3SS was induced by shifting to 37°C and depleting Ca^2+^ for 3h.

In our previous study, the samples subjected to transcriptome sequencing (RNA-Seq) had been collected after 3 hours under virulence-inducing conditions. To rule out the possibility that the low expression of T3SS genes observed in the ∆*rpoN* strain was a cumulative indirect effect of RpoN absence, we performed RNA-Seq on WT and ∆*rpoN* bacteria grown under virulence-inducing conditions for different periods of time. Already, at 15 or 30 minutes after inducing virulence, the mRNA levels of several T3SS genes, especially *ysc* genes, were lower in the ∆*rpoN* strain compared with the WT strain (Fig. 2). This difference in mRNA abundance between the strains increased at 90 minutes post inducement and was even more pronounced at 180 minutes post inducement (Fig. 2 and Fig. 1A).

**Fig 2.**
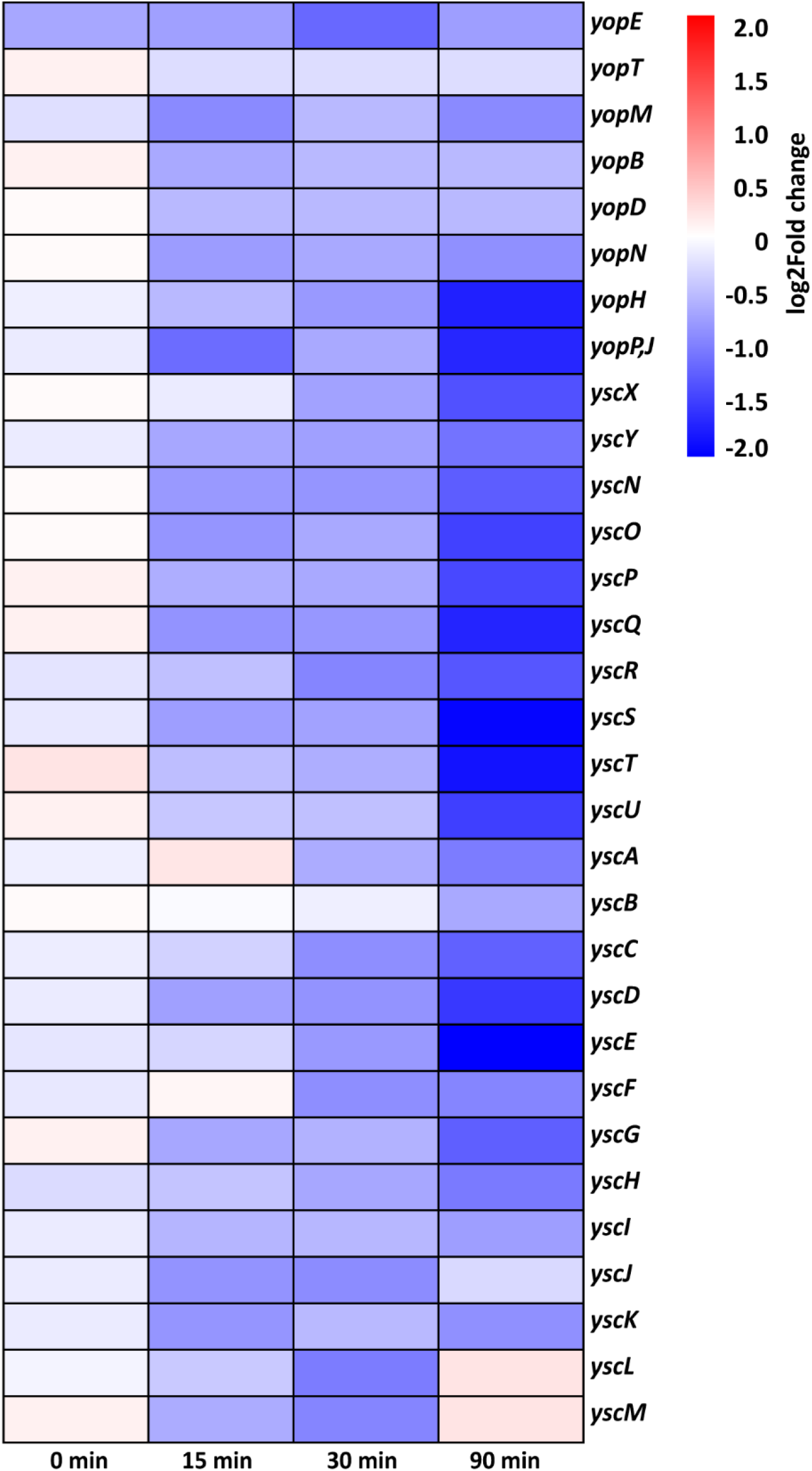
Absence of RpoN affects early gene expression associated with inducement of T3S. Heat map showing mRNA expression profiles of T3SS structural component (*ysc*) genes and *Yersinia* outer protein genes in ∆*rpoN* compared with the WT strain. Relative gene expression is shown for 0, 15, 30 and 90 minutes of virulence plasmid inducement (37°C and depletion of Ca^2+^). Colour scales are based on relative expression in log2 fold change. Blue indicates downregulation in ∆*rpoN*.

### A putative role of the enhancer-binding protein GlrR in RpoN-mediated activation of T3S

Since RpoN has been shown to regulate the T3SS of different plant pathogens with the help of specific EBPs ^24-27^, we scrutinised the *Y. pseudotuberculosis* genome to identify hypothetical EBPs. We identified seven hypothetical EBPs (Table S1) and generated a deletion mutant for each corresponding gene. Then, we tested T3S in these mutants and found that one of them, the ∆*glrR* strain, showed weaker secretion than the WT strain (Fig. 3). The decrease in T3S was less pronounced compared to that seen for the ∆*rpoN* strain. Nevertheless, this result suggests that GlrR-mediated activation of RpoN contributes, at least partially, to inducing T3S.

**Fig 3.**
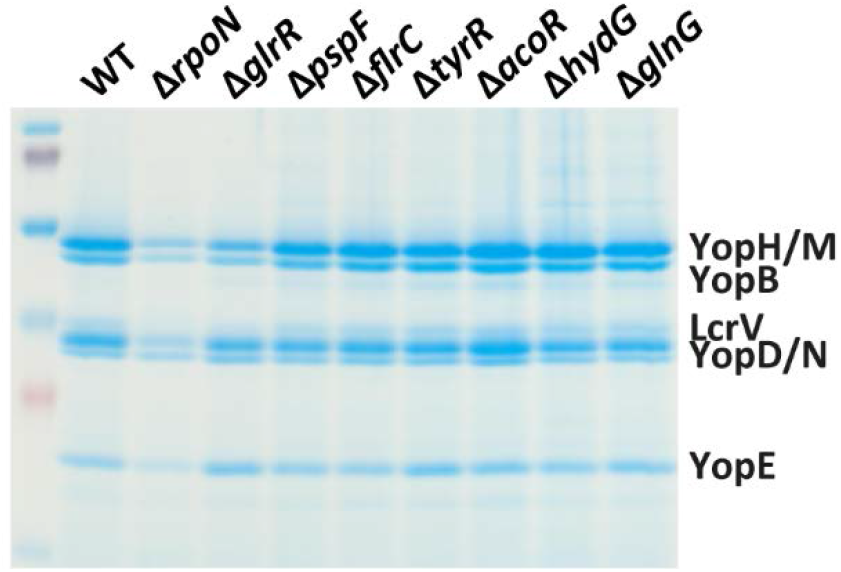
Deletion of the gene encoding the EBP GlrR results in reduced secretion of Yop effectors. T3S profile of WT *Y. pseudotuberculosis*, the isogenic mutants ∆*rpoN* and seven different EBP mutants (∆*glrR*, ∆*pspF*, ∆*flrC*, ∆*tyrR*, ∆*acoR*, ∆*hyd*G and ∆*glnG*). T3SS was induced by shifting to 37°C and depleting Ca^2+^ for 2h.

### No binding sites for RpoN on the *Yersinia pseudotuberculosis* virulence plasmid

To investigate the possibility of a direct effect of RpoN on the transcription of genes on the virulence plasmid, we re-analysed our ChIP-Seq data reported earlier ^9^ using a more relaxed cut-off of 1.7-fold difference over genomic noise instead of 2.5-fold and screened for putative RpoN binding sites located on the virulence plasmid. This analysis identified three putative binding sites (BSs) in the region of the beginning of the *yscN-U* operon, two upstream of *yscN* and one inside the *yscN* open reading frame (Fig. S1). All these were mutated, and mutation of the two binding sites upstream of yscN did not reduce the T3S, but two out of three different mutations of the putative RpoN-binding site located in *yscN*, upstream of the yscO gene, did result in reduced T3S (Fig. S1). For verification of RpoN-binding to these sites, an EMSA assay for the potential binding sites was employed. However, this assay revealed nothing that pointed to a productive RpoN binding to this region. (Fig. S1). Hence, the reduction in T3S is not caused by the loss of a direct binding of RpoN to DNA sequences on the virulence plasmid; rather, this is an indirect effect caused by the absence of RpoN.

### T3S-correlated inducement of *rpoE* is abolished in the absence of *rpoN*

Indirect effects of deleting global regulators, such as sigma factors, are common where some genes are induced as a stress response to compensate for the loss of the alternative sigma factor. Examples here are the inducement of the phage shock protein operon to compensate for the loss of RpoE in *Salmonella* ^*35*^. There are also many examples of hierarchical regulatory networks, where regulators control other regulators, and where the loss of the former leads to downstream effects attributed to the latter. This is also true for sigma factors, where for example expression of RpoH has been reported to be regulated by RpoE in *E. coli* ^36^. A similar scenario was also obvious in our previous study, where we used ChIP-seq and RNA-seq to determine the RpoN regulon in *Y. pseudotuberculosis*. The deletion of *rpoN* was associated with altered expression of genes encoding global regulators, among them *rpoD, rpoE* and *rpoS* ^9^. Three hours after inducing T3S, the expression of *rpoD* was significantly higher, whereas the expression of *rpoE* was much lower both in the stationary phase and under virulence-inducing conditions in ∆*rpoN* compared to that in the WT strain. On the other hand, *rpoS* was higher in the stationary phase but lower under inducement conditions. To investigate involvement of different sigma factors in inducing T3S, we first analysed their expression at 15, 30 and 90 min after T3S inducement in both the WT and the ∆*rpoN* strain. Interestingly, in the WT strain there was a clear shift during the period of inducement, with a decrease in levels of *rpoD* mRNA and a concomitant increase in *rpoE* mRNA and also an early but less pronounced increase in *rpoS* mRNA (Fig. 4A). The amount of *rpoN* mRNA was unchanged, and *fliA* mRNA was decreased. Interestingly, almost the opposite was seen in the ∆*rpoN* strain, except for *fliA*, which was downregulated as in WT (Fig. 4A). Hence, the sigma factor regulatory networks are clearly affected in the absence of RpoN. The lack of an increase in RpoE, and maybe also of RpoS upon initiation of T3S, might be responsible for the observed reduction in transcription of T3S genes.

**Fig 4.**
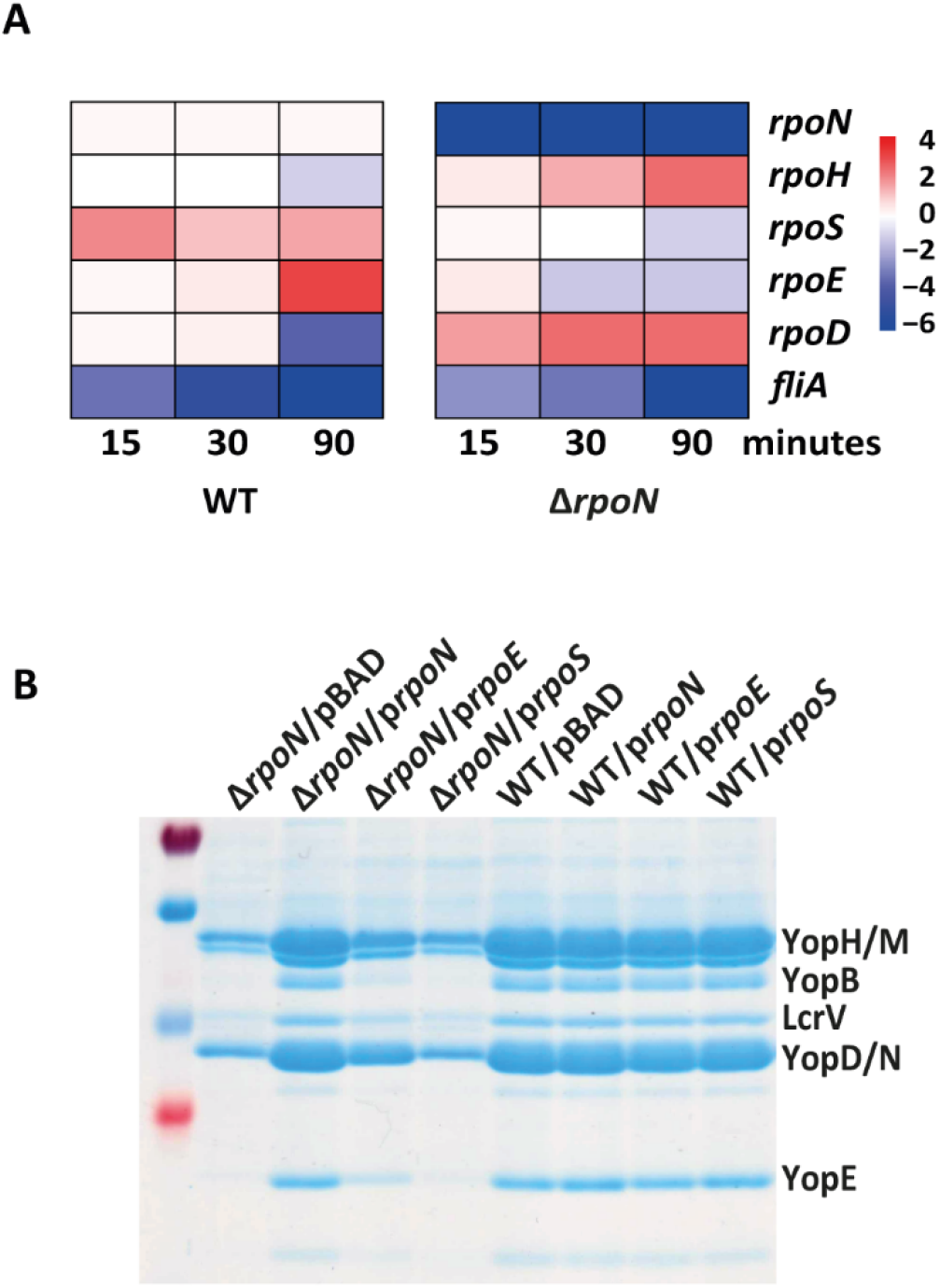
Expression of *rpoE* is increased during T3SS inducement in WT but not in ∆*rpoN*, and over-expression of RpoE partially restores the T3S defect in ∆*rpoN*. (A) Heat map showing time course expression profiles of sigma factor encoding genes in WT *Y. pseudotuberculosis*, and the isogenic ∆*rpoN* strain. Gene expression at 15 minutes, 30 minutes, and 90 minutes of plasmid inducement (Ca^2+^ depletion) relative to uninduced (0 min) is shown. Colour scale is based on relative expression in log2 fold change. Blue indicates downregulation in the ∆*rpoN* strain. (B) T3S profiles of WT and ∆*rpoN* strains overexpressing in *trans* the sigma factors RpoN, RpoE and RpoS, and of control empty-vector (pBAD).

Given the shift in expression from RpoD to RpoE upon inducing T3S, we also checked for potential RpoE binding sites in the promoter regions of differentially expressed genes in the WT strain 90 minutes after T3S inducement, where RpoE expression is much higher compared to time point zero 0 (Fig. 4A). We found that 40% of the differently expressed genes, 600 out of 1520, have potential RpoE binding sites. However, as RpoE binding motifs are poorly conserved ^37^, the *in silico* efficacy may be low and could predict some false positives or fail to predict true biologically active binding sites. However, RpoE binding sites present in high numbers among differentially expressed genes and higher expression of *rpoE* in corresponding conditions seem to indicate a role for RpoE.

### A potential role for RpoE in inducing T3S

To investigate a possible importance of RpoS and RpoE for inducement of T3S in *Y. pseudotuberculosis*, we next set out to construct deletion mutants of *rpoS* and *rpoE* to determine the eventual effects on T3S in the resulting mutant strains. However, the *rpoE* gene turned out to be temperature sensitive in *Y. pseudotuberculosis*, and thereby not able to be induced for T3S, which is in accordance with what reported earlier ^38^. The resulting Δ*rpoS* strain showed similar levels of T3S as the WT strain, thus excluding this sigma factor as being important in the RpoN regulatory network leading to a functional T3SS.

As an alternative, we also overexpressed the genes encoding the sigma factors in *trans* in ∆*rpoN* and subjected the resulting strains to T3S assays. Here we found that *rpoE*, but not *rpoS*, overexpression partially restored the WT T3S positive phenotype (Fig. 4B). This result, together with the observed lack of upregulation of this gene during virulence-inducing conditions, suggests that the T3S defect of the ∆*rpoN* strain might be at least partially due to mis-regulation of RpoE.

Given the potential role of RpoE in regulating virulence inducement in *Y. pseudotuberculosis*, we next performed an *in silico* prediction of binding sites on the *Y. pseudotuberculosis* virulence plasmid. This analysis revealed potential binding sites, especially for RpoE, RpoS and RpoD, upstream of the main *ysc* operons that were found downregulated in the ∆*rpoN* strain (Fig. 5). Hence, together with the observed partial rescue of T3S in the ∆*rpoN* strain by overexpressing RpoE, the presence of RpoE binding sites upstream of the affected operons supports the suggestion of a role for this sigma factor in the mechanism mediating inducement of T3S.

**Fig 5.**
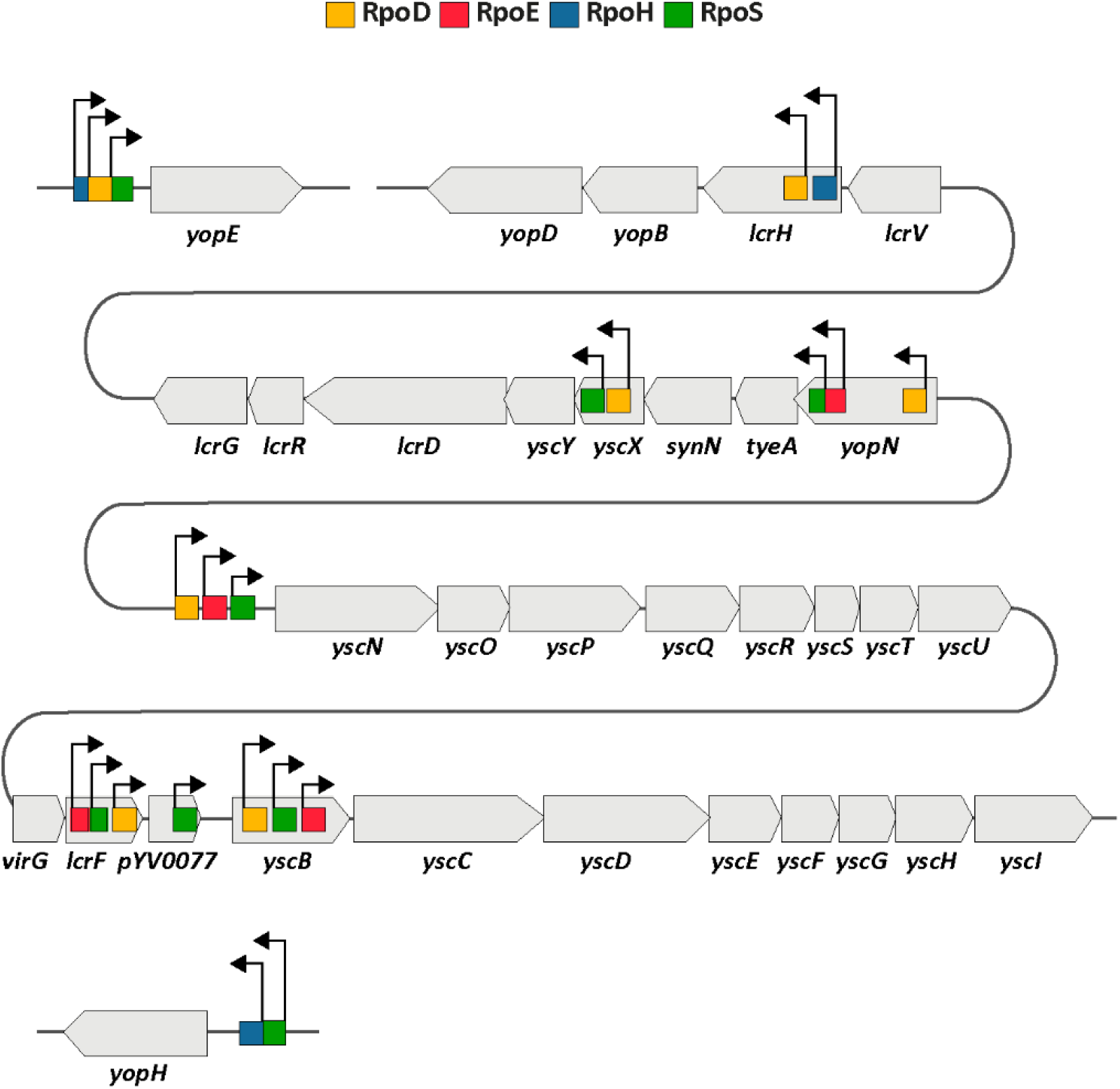
Potential sigma factor cross talk controlling T3SS gene expression. *In silico* prediction of sigma factor binding sites at intra- or intergenic regions of associated genes. Arrow signs indicate the direction of sigma factor binding motifs. Motif lengths are not scaled according to gene length or intergenic distance.

## Discussion

We recently showed that the alternative sigma factor RpoN is essential for the virulence of the intestinal pathogen *Y. pseudotuberculosis*. Therefore, in this study we investigated the probable role of RpoN in regulating the for virulence essential T3SS, which allows this pathogen to survive extra-cellularly in lymph nodes that abounds in immune cells. Data presented indicate that the requirement of this sigma factor for *Y. pseudotuberculosis* virulence involves its positive effect on the T3SS. There is a severe reduction in T3S in strains lacking the *rpoN* gene, and our data imply that this is due to an insufficient transcription of the genes encoding T3SS structural components. The mechanism responsible for this appears to partly involve diminished expression of RpoE occurring down-stream of RpoN.

Our data commonly points towards a more global role for RpoN in the regulatory networks involved in mounting full virulence of *Y. pseudotuberculosis*. We have not found any evidence of a direct effect of RpoN binding to the virulence plasmid, either in the previous ChIP-Seq study or by evaluating possible binding sites predicted with less stringency by mutational studies and EMSA assays (Fig S1). However, in contrast to that of RpoN, there were clear indications of binding sites for other sigma factors on the plasmid, sequences of high homology to reported binding regions for RpoE, but also for RpoS and RpoD. Whether these sites are functional remains to be evaluated in detail. A role for RpoE in regulation of *Y. pseudotuberculosis* T3SS has indeed been suggested previously ^39^, as for T3SS in *E. coli* and *Salmonella* ^39^.

Bacteria can meet environmental challenges by involving diverse signalling cascades that can merge into a common stress response. It is therefore not surprising if the inducement of T3S, which involves sensing with subsequent activation of both metabolic and structural functions, requires gene products of diverse regulons. The RpoN regulon might mediate activation of another regulator, such as RpoE in this case, and there might also be additional hierarchical levels of transcriptional regulation. This type of sigma factor cooperation has for example been shown for flagellar biosynthesis in *P. aeruginosa* with a four-level hierarchy of regulators ^40^. Other examples are sigma factor cascades in Salmonella, where RpoE and RpoH promote translation of RpoS by increasing the expression of Hfq ^35^.

For inducing T3S, our transcriptomic data indicate a shift in sigma factor usage, from RpoD to RpoE, and to some extent RpoS. This shift correlates with the environmental cues encountered by the bacteria, with a drastic increase in temperature from 26°C to 37°C and a simultaneous change in extracellular ion composition, constituting both a sudden initial stress for RpoS to cope with and extra-cytoplasmic changes for RpoE. The scenario may involve an increased formation especially of RpoE holoenzymes and concomitant transcription instead of that of the house keeping sigma factor RpoD. In addition, it is likely that the growth inhibition that occurs about 45–60 minutes after T3S inducement leads to increased availability of core RNAPs due to reduced transcription of rRNA ^41,42^, as suggested for the stringent response ^43,44^. The partial rescue of T3S in the *rpoN* mutant strain by overexpression of RpoE supports a role for this sigma factor in regulating the formation of the T3SS apparatus, which is in agreement with other studies proposing a link between T3S and extra-cytoplasmic responses in *Y. pseudotuberculosis* ^39^. This is further supported by the identification of RpoE binding sites upstream of the operons on the virulence plasmid that were downregulated in the *rpoN* mutant strain. Hence, taken together, our data hints to a potential direct effect of RpoE, which has to be elucidated further at a molecular level. Since long it has been known that the RpoE homolog HrpL mediates activation of T3SS in plants ^45^, and interestingly, a similar regulatory path involving RpoN and RpoE as we suggest for inducing T3S in *Y. pseudotuberculosis* has been reported for the plant pathogen *Erwinia amylovora*, where RpoN acts upstream of the RpoE homolog HrpL ^22^. There is also a recent study by Shao and co-workers (2021) showing that several T3SS genes in *Pseudomonas syringae* could be targets of HrpL ^46^.

The finding of a likely role of the RpoN/GlrR regulon in inducing productive T3S in *Y. pseudotuberculosis* is interesting, and even if the RpoN/GlrR regulon in *Y. pseudotuberculosis* remains to be determined, this finding further supports an important upstream role of RpoN. The EBP GlrR is the response regulator of the two-component system GlrR/GlrK (also named QseF/QseE or YfhA/YfhK) that contributes to cell envelope integrity by regulating transcription of the sRNAs GlmY ^47^. RpoN/GlrR-mediated control of GlmY and the neighbouring sRNA GlmZ is conserved in many enterobacteria ^48^. GlmY is a central player in a complex regulatory cascade controlling production of glucosamine-6-phosphate that initiates cell envelope biosynthesis ^49^. Also interesting is that this GlrR has been shown to regulate expression of RpoE in *E. coli*. Hence, GlrR and RpoE might act in the same RpoN pathway in regulating T3S. However, the observed contribution of RpoE and also of GlrR is only partial, so there might be other RpoN downstream effects that contribute to T3S inducement.

Taken together, our data imply an important role for RpoN in mediating efficient inducement of T3S in *Y. pseudotuberculosis*. We provide data pointing to possible explanations behind the observed requirement of RpoN for a functional T3SS in *Y. pseudotuberculosis*, and thereby its ability to cause disease. This involves an RpoN-mediated regulatory network with downstream activation of RpoE, possibly via the EBP GlrR.

## Materials and methods

### Bacterial strains and growth conditions

Strains and plasmids are listed in Table S2 in the supplemental material. *Yersinia pseudotuberculosis* strain YPIII was used in this study. *Escherichia coli* S17-1 λpir was used for cloning and conjugation. Antibiotics were used at the following concentrations: ampicillin (100 µg/ml), kanamycin (50 µg/ml) and chloramphenicol (25 µg/ml). *Y. pseudotuberculosis* and *E. coli* strains were routinely grown in Luria-Bertani (LB) broth or agar at 26°C and 37°C, respectively. For virulence inducement, overnight bacterial cultures were diluted to an OD600 of 0.05 in LB and grown at 26°C. After 1 hour, calcium was depleted by adding 5 mM EGTA and 20 mM MgCl2 and the cultures were shifted to 37°C ^50^.

### Construction of mutant strains

In-frame gene deletion and binding-site mutations in *Y. pseudotuberculosis* were performed using an In-Fusion HD cloning kit (Clontech) according to the manufacturer’s instructions. Briefly, the flanking regions of the respective gene were amplified by PCR and cloned into the suicide vector pDM4. This construct was used to transform S17-1 λpir and then transferred into the recipient strain through conjugation. Transconjugants were purified and incubated on 5% sucrose to recombine out the vector together with WT sequence. Deletion or mutation was confirmed by PCR. For trans-complementation, the gene was PCR amplified and cloned into the pBAD24 plasmid. All constructs were verified by sequencing.

### Analysis of Type III protein secretion by Y. pseudotuberculosis

Overnight Y. pseudotuberculosis cultures were diluted to an OD600 of 0.05 in LB and allowed to grow for 1 hour at 26°C. Cultures were then shifted to 37°C and grown under T3S-inducive conditions (5 mM EGTA and 20 mM MgCl2). Aliquots were removed at 2 or 3 hours post inducement. Samples containing secreted proteins were prepared from bacteria-free supernatants recovered after centrifugation. The secreted proteins were precipitated with 10% TCA and separated by 12% SDS-PAGE.

### Western blot detection of YscN

For detection of YscN in the different strains analysed, the T3SS was induced as described above. After 3 hours of growth under T3S-inducing conditions, cells were pelleted by centrifugation, the supernatant discarded, and the cell pellet resuspended in Laemmli sample buffer and boiled for 5 min. Samples were loaded according to the OD600 values of cultures and separated by 12% SDS-PAGE, transferred to an Immobilon-P PVDF membrane (Millipore) and probed overnight at 4°C with a primary rabbit anti-YscN antibody at a 1:2000 dilution. The final blot was developed by chemiluminescence (Immobilon Western Chemiluminescent HRP Substrate, Millipore) and the resulting bands imaged using an Amersham Imager 680.

### cDNA preparation and qPCR

To validate the RNA-Seq results, qPCR was performed using KAPA SYBR FAST qPCR Master Mix (KAPA Biosystems) and a Real-Time PCR Detection System (Bio-Rad). Bacterial strains were grown for 2 hours at 37°C under inducing conditions in LB, and the total RNA was extracted as described below. Isolated RNA was used as the template for cDNA synthesis using a RevertAid First-Strand cDNA synthesis kit (Thermo Scientific). Experiments were done in triplicate. Six plasmid genes were tested, and the stable reference genes YPK_0316 and YPK_1676 were selected as internal controls to calculate the relative expression levels of the tested genes using appropriate primers (see Table S3 in the supplemental material).

### RNA isolation and Illumina sequencing

The total RNA was isolated from three independent biological replicates of WT *Y. pseudotuberculosis* and an isogenic *rpoN* deletion strain under virulence-inducing conditions at different time points. Specifically, cells were grown exponentially at 26°C, shifted to 37°C and incubated in a Ca^2+^-depleted medium. Samples for RNA isolation were taken at 0, 15, 30 and 90 minutes after inducement. RNA isolation, purification and treatment, preparation of cDNA libraries, Illumina sequencing and quality check, and processing of raw reads were done as described in our previous study ^9^. ProkSeq, a complete RNA-Seq data analysis package for prokaryotes, was used for further RNA-Seq data processing, quality control and visualisation ^51^. Reads were aligned to the *Y. pseudotuberculosis* plasmid (NC_006153) using Bowtie 2 ^52^ with the unique-mapping option. Post-mapped read quality checking was done using RSeQC ^53^, and the number of reads for each gene was counted using featureCounts ^54^. Differential gene expression was determined using DESeq2 and by using the shrinkage estimation of the dispersion option (lfcSrink =True) to generate more accurate estimates of differential expression in fold changes ^55^. Differential expression of a gene was defined using absolute log2 fold change values ≥ 1.5 and a false-discovery rate value of 0.05. Figures were plotted using the ggplot2 package in R (Linux version 4.0.2) and GraphPad Prism (version 8.0).

### ChIP-Seq data analysis

ChIP-Seq data analysis, data visualisation, downstream bioinformatic and computational analyses, and quality check and processing of raw reads were done as described in our previous study [9]. Peak calling was done using MACS (2.1.2) ^56^ with the following custom settings: log2FC ˃ 1 over input, –broad –g, –broad-cutoff = 0.2. Identified peak coordinates in the genome were used to identify the probable regulated genes using the R packages. A position weight matrix (PWM) of the predicted motif was used to map the motif presence around the peak centre using the FIMO-meme package (Linux version) ^57^. A region of 50 bases upstream and downstream of the peak centre was used as input to search for the PWM of the motif.

### Electrophoretic Mobility Shift Assay (EMSA)

The 40-ng samples of different DNA probes (350-nt genomic regions of *yscN* and *pspA*) were mixed with various amounts of purified RpoN protein in 20 μl of the sample buffer, (50 mM HEPES, pH 7.9, 0.1 mM EDTA, 100 mM NaCl, 2.8% polyethylene glycol 8000, 100 mM KCl, 1.0 mM dithiothreitol, 10 mM MgCl2, 5% glycerol, and 100 μg/ml bovine serum albumin). The reactions were incubated for 20 min at room temperature. To analyse the samples, 6% native polyacrylamide gel containing 2.5% glycerol and 0.5× Tris-borate-EDTA (TBE) buffer was used. Gels were run at 90 V for 120 min and thereafter stained with Ethidium bromide at room temperature for 5 min and visualised using a Gel Doc XR imaging system (Bio-Rad).

## Data availability

The RNA-seq data files have been deposited in Gene Expression Omnibus under accession number GSE195976. All the computer code and pipeline used in these studies are available on request.

## Acknowledgements

The work has been supported by funding from Knut and Alice Wallenberg foundation (2016.0063), Swedish research Council (2018-02855), and the Medical faculty at Umea University. We thank SciLife for sequencing facilities.

## Contributions

Conception or design of the work: A.K.M.F.M, R.N, K.N, R.C and M.F; data collection: A.K.M.F.M, K.N, R.N, data analysis and interpretation: F.A.K.M.M, R.N, K.N, and M.F.; drafting of the article: F.A.K.M.M, R.N, K.N and M.F; critical revision and contributions to the article: A.K.M.F.M, R.N, K.N, R.C, K.A. and M.F.; final approval of the version to be published: A.K.M.F.M, K.N,R.N, R.C, K.A. and M.F.

## Figure legends

**Fig S1.**
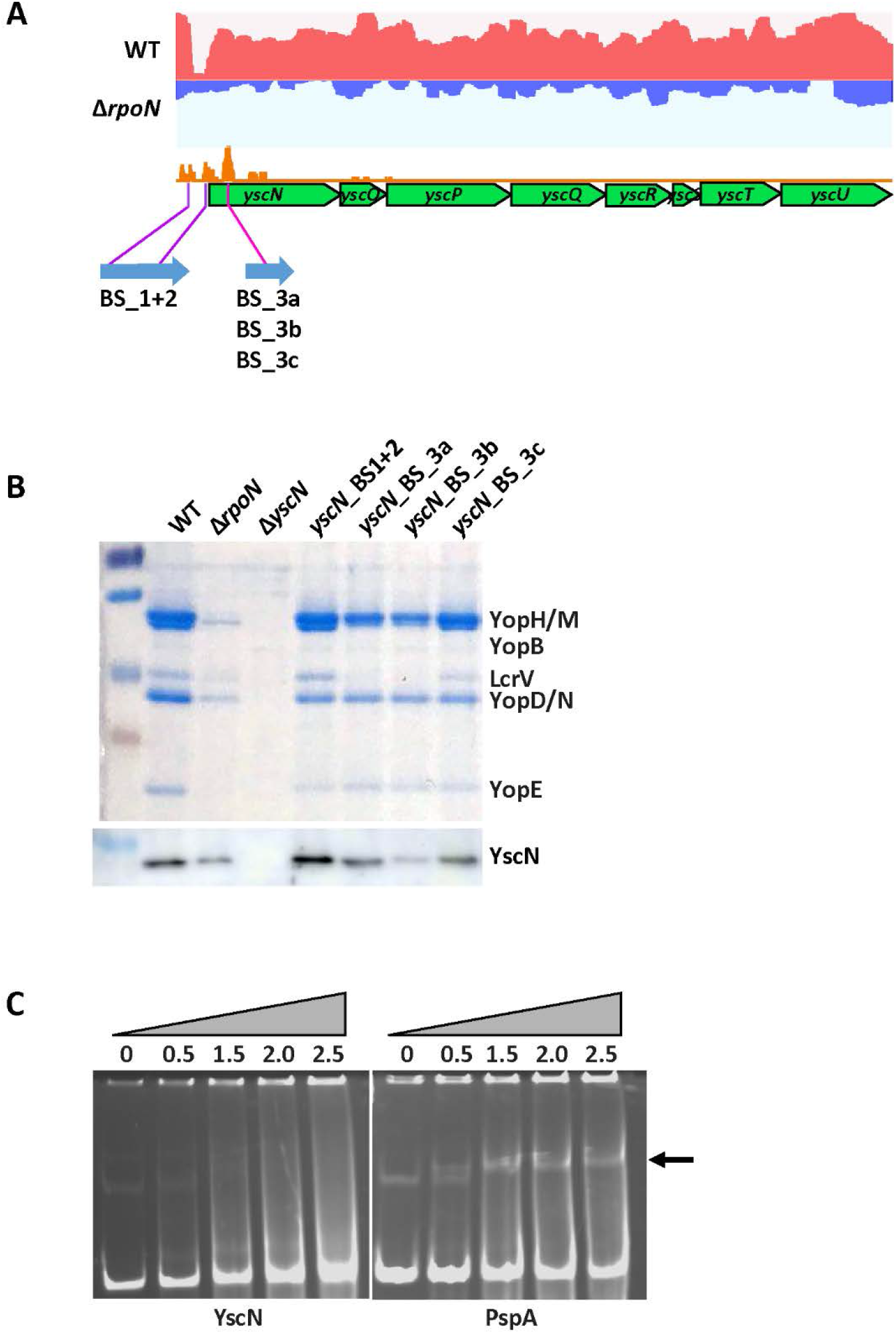
Mutations in the putative RpoN binding site located in gene *yscN* negatively affect T3S in *Y. pseudotuberculosis*. (A) Putative RpoN binding sites located upstream of *yscN* and in its open reading frame. Red and blue track indicates the RNA-Seq read coverage of the *yscN-*U operon in the WT and ∆*rpoN* strains at 90 minutes after inducement. The read coverage tracks are normalised according to library depth (B) The upper part shows the T3S profiles of WT *Y. pseudotuberculosis*, ∆*rpoN* and strains carrying mutations in putative RpoN binding sites. The lower part shows a western blot analysis of YscN in the mentioned strains. (C) EMSA of RpoN binding to potential RpoN-binding sites in *yscN*. An RpoN-binding site located upstream of *pspA* was included as a positive control. Increasing concentrations of the RpoN protein were mixed with 350-bp fragments covering potential RpoN-binding sites. Arrow indicates the expected band of RpoN-DNA complex.

---

**Table S1:** Genes encoding putative RpoN enhancer-binding proteins present in *Y. pseudotuberculosis* YPIII.

| Gene name<br>(KEGG* accession number) | Definition | KEGG Orthology |
| --- | --- | --- |
| *glrR* (YPK_1259) | Two-component, sigma54 specific, transcriptional regulator, Fis family | Two-component system, NtrC family, response regulator GlrR |
| *pspF* (YPK_1893) | Sigma54 specific transcriptional regulator, Fis family | *psp* operon transcriptional activator |
| *flrC* (YPK_0709) | Sigma54 specific transcriptional regulator, Fis family | Two-component system, response regulator FlrC |
| *tyrR* (YPK_1902) | Transcriptional regulator, TyrR | Transcriptional regulator of *aroF*, *aroG*, *tyrA* and aromatic amino acid transport |
| *acoR* (YPK_1975) | GAF modulated sigma54 specific transcriptional regulator, Fis family | Sigma-54 dependent transcriptional regulator, acetoin dehydrogenase operon transcriptional activator AcoR |
| *hydG* (YPK_2907) | Two-component, sigma54 specific, transcriptional regulator, Fis family | Two-component system, NtrC family, response regulator HydG |
| *glnG* (YPK_4191) | Nitrogen metabolism transcriptional regulator, NtrC, Fis Family | Two-component system, NtrC family, nitrogen regulation response regulator GlnG |

\* KEGG: Kyoto Encyclopedia of Genes and Genomes.

**Table S2:**
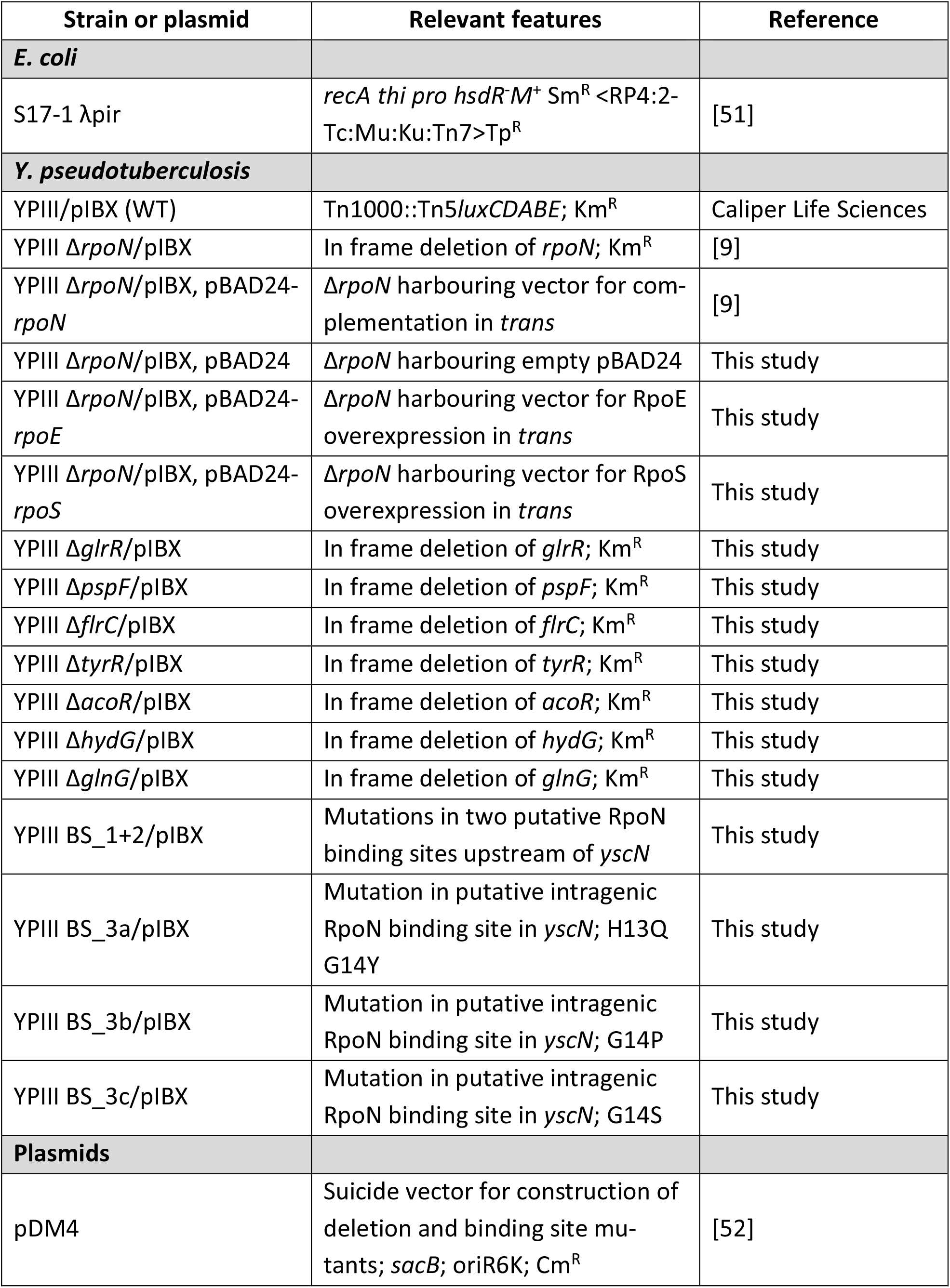
Bacterial strains and plasmids used in this study.

**Table S3:**
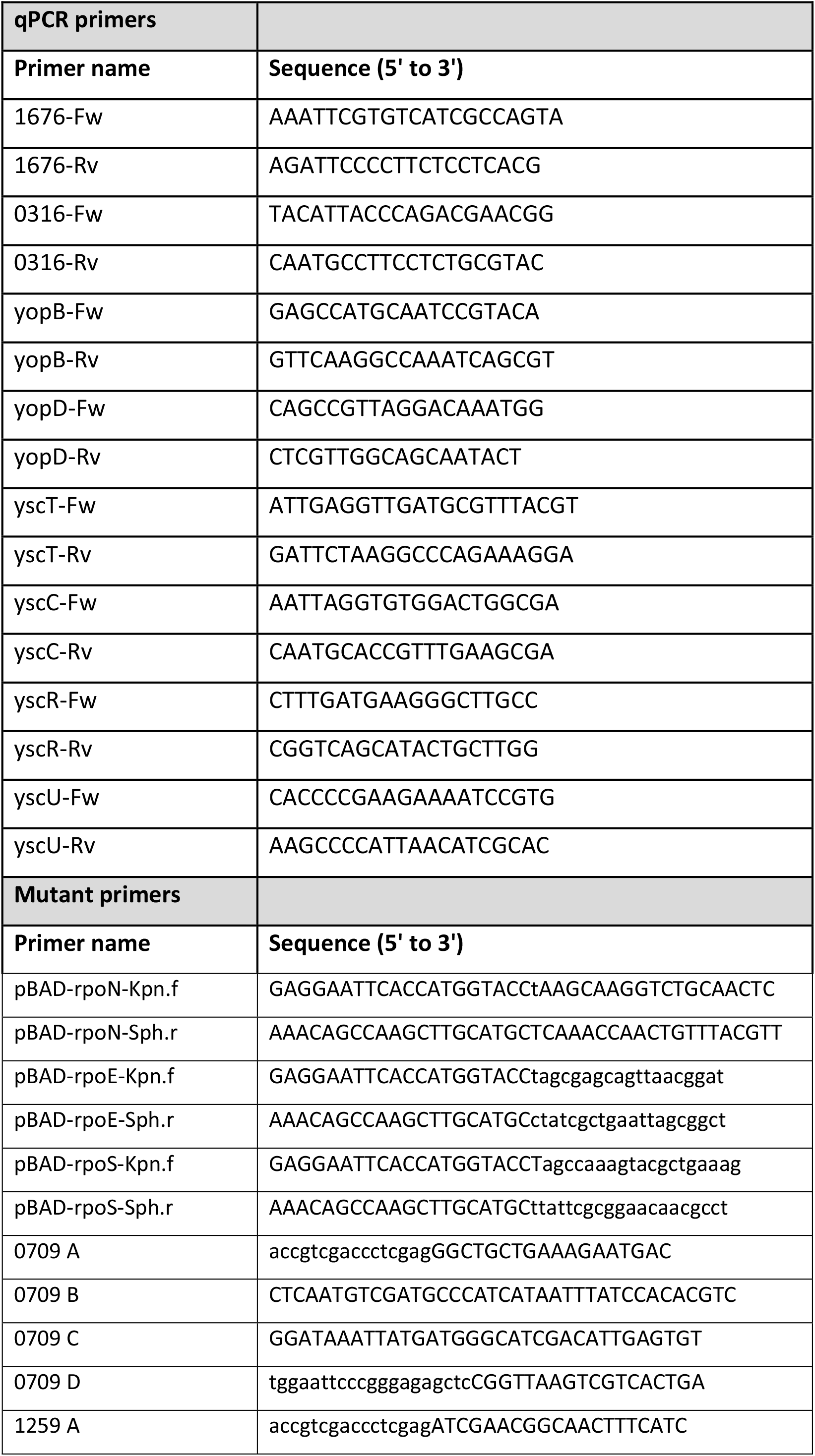
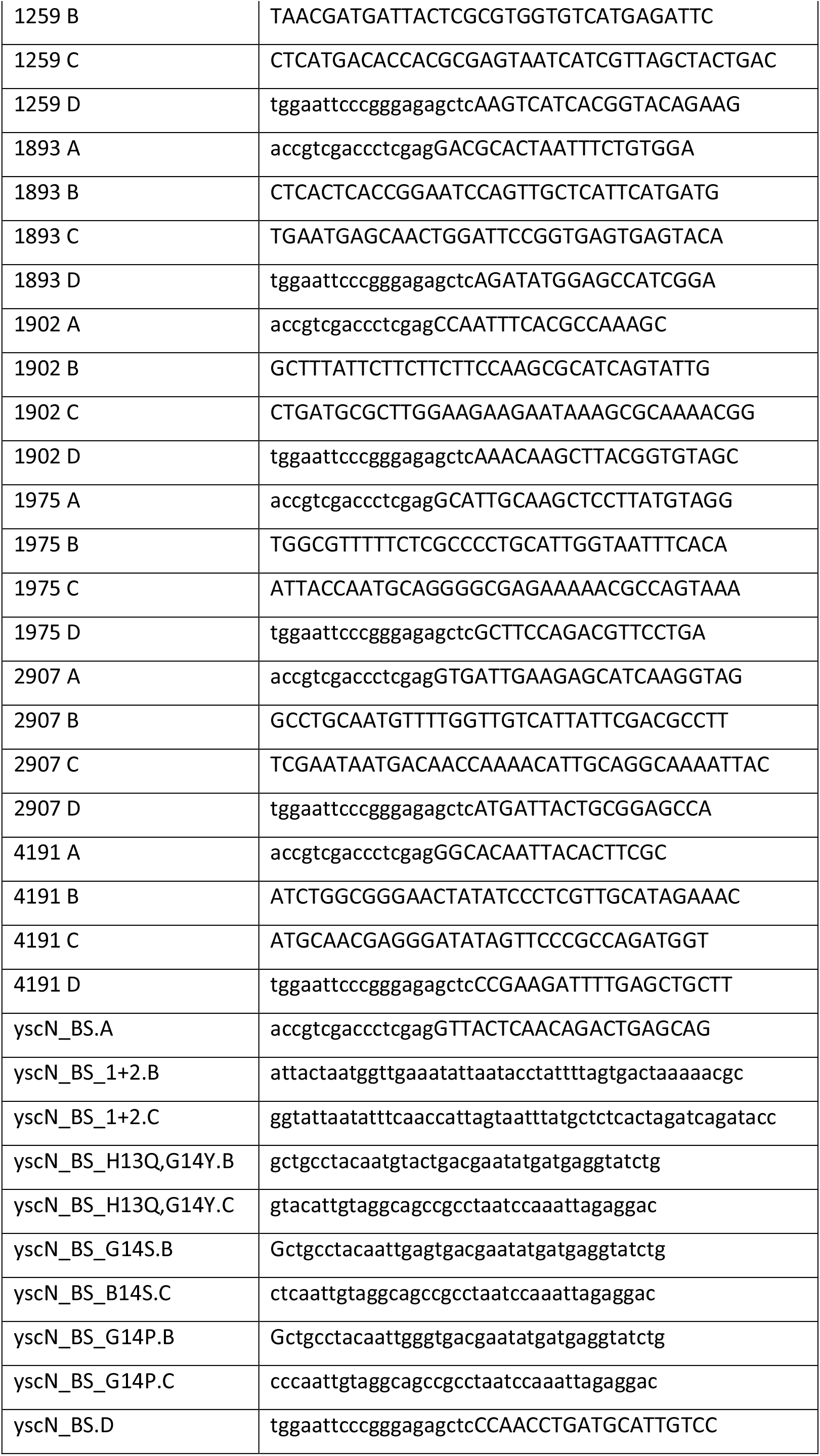
Oligonucleotide primers used in this study.

